# Strengthening marine amphipod DNA barcode libraries for environmental monitoring

**DOI:** 10.1101/2020.08.26.268896

**Authors:** Chinnamani Prasannakumar, Ganesh Manikantan, J. Vijaylaxmi, Balakrishnan Gunalan, Seerangan Manokaran, S. R. Pugazhvendan

**Affiliations:** Biological Oceanography Division, CSIR-National Institute of Oceanography, Dona Paula, Panaji, Goa-403004, India; Institute of Marine Microbes and Ecosphere, State Key Laboratory for Marine Environmental Sciences, Xiamen University, Xiamen, Fujian, 361102, PR China; Centre of Advance studies in Marine Biology, Annamalai University, Parangipettai, Tamil Nadu- 608502, India; Department of Marine Sciences, Goa University, Taleigao Plateau, Goa-403206, India; Post Graduate and Research Department of Zoology, Thiru Kolanjiappar Government Arts College, Virudhachalam, Tamil Nadu- 606001, India; Center for Environment & Water, King Fahd University of Petroleum and Minerals, Dhahran-31261, Saudi Arabia; Department of Zoology, Arignar Anna Government Arts College, Cheyyar, Tamil Nadu- 604407, India; Department of Zoology, Annamalai University, Annamalai Nagar, Chidambaram, Tamil Nadu- 608002, India

**Keywords:** Marine amphipods, Environmental monitoring, COI, DNA barcoding, Amphipod barcoding

## Abstract

Environmental DNA barcoding technology is gaining innovative applications. The effectiveness of current DNA barcode reference libraries in identifying amphipod barcodes and/or strengthening the existing library was tested. From 2500 amphipod individuals we barcoded 22 amphipod species belonging to 17 genera, 13 families among which 13 species were first time barcoded. More than 80 percent of the species were new distributional records. The minimum and maximum inter-specific pair-wise distance values was respectively 0.16 and 5.51 percent. Defining family specific species threshold values would be imperative, rather than expecting a universal barcode gap for amphipod species. The overall mean pair-wise distance, nucleotide diversity and Tajima’s statistics were 3.59 percent, 0.27 and 2.62, respectively. There is a strong need to increase the number of amphipod species barcodes in the reference database. For better facilitation of environmental monitoring, the datasets could be exclusively accessed at BOLD through http://dx.doi.org/10.5883/DS-MAOI.

## 1. Introduction

Amphipods (Phylum: Arthropoda, Class: Malacostraca, Order: Amphipoda) are a significant invertebrate fauna associated with coastal ocean environments which connects producers and consumers (such as fishes) in marine trophic webs (Sanchez-Jerez et al. 1999; Zakhama-Sraieb et al. 2006; Fernandez-Gonzalez and Sanchez-Jerez 2014). Amphipods have been used as key species in the assessment of environmental quality because they live in close proximity to marine and estuarine sediments (Chapman et al., 1992, Chapman et al., 2013, Postma et al., 2002) and are used as a bio-indicator of contamination since they are responsive to changes in environmental conditions (Bellan-Santini 1980; Virnstein 1987; Conradi et al. 1997; Guerra-Garcia and Garcia-Gomez 2001). Example; oil spills affects growth and abundance of amphipod fauna (Gesteira et al., 2000; Andrade & Renaud, 2011; Joydas et al., 2012; Lotufo et al., 2016) and the bioaccumulation of hydrocarbons differs between families of amphipods (Lourenco et al., 2019). Amphipods are used to monitoring sediments acute toxicity which was proven to be species-specific (Ohji et al., 2002; Vacchi et al., 2019). Amphipods were also used to biomonitor the trace metal concentrations in coastal environments (Rainbow et al., 1998; Fialkowski et al., 2009; Morrison et al., 2017). By shredding plastic carrier bags, amphipod species were known to generate numerous pieces of microplastics which in turn significantly reduces amphipod’s algal consumption (Hodgson et al., 2018; Carrasco et al., 2019). Changes in algal nutritional quality due to ocean acidification was also known to modify amphipod’s feeding behavior (Benítez et al., 2016). Other environmental parameters such as light intensity and salinity influences the growth and the composition of amphipod assemblages (Navarro-Barranco & Hughes, 2015; de-la-Ossa-Carretero et al., 2016). Therefore, the composition of the amphipod species typically represents the environmental conditions where they have been isolated.

Conventional taxonomy tussles to classify amphipods as they were small sized with poor taxonomic descriptions and converging morphological characters (Knowlton, 1993, Radulovici et al., 2010), making them an ideal group for DNA barcoding application. DNA barcoding involves sequencing a gene fragment from precisely identified specimens to form a database and enabling species identification (even by non-experts) by comparing the same gene sequences sequenced from unidentified specimens (Hebert et al., 2003, Mitchell, 2008). DNA barcode based identification was successful in marine amphipods of Arctic (Tempestini et al., 2018), Antarctic (Havermans et al., 2011), Atlantic (Costa et al., 2009) and Pacific (Jażdżewska and Mamos, 2019) Oceans. Such efforts, however are rare in Indian Ocean realms where amphipod diversity were relatively richer (Mondal et al., 2010, Raja et al., 2013).

Environmental DNA (eDNA) is the DNA recovered from environmental substances such as water, soil, or sediment (Taberlet et al. 2012; Thomsen and Willerslev 2015) and eDNA barcoding refers to sequencing the DNA barcodes, amplified from DNA recovered in the environmental for discovering the taxonomy and biodiversity of the sampled area. eDNA barcoding technology is finding innovative applications for monitoring marine biodiversity (Nguyen et al., 2020), from deep sea hydrothermal vents (Cowart et al., 2020) to plankton gut materials (Oh et al., 2020) and also in tracking terrestrial biodiversity (Heyde et al., 2020). eDNA barcoding technology has also been shown to be useful in monitoring the marine ecosystem to detect invasive species (Kim et al., 2020) even from the DNA recovered from marine litters (Ibabe et al., 2020).

eDNA technology has been shown to be useful in spatial and temporal monitoring in a wide variety of settings, as the methods are relatively inexpensive, reliable and quicker than conventional monitoring (Lecaudey et al. 2019; Preissler et al. 2019; Reinhardt et al. 2019; Sutter and Kinziger 2019; Sales et al. 2020). The breakthrough in eDNA barcoding technology is in its ability to monitor the ecosystem without causing unnecessary harm to ecosystem or their organisms by non-invasive sampling strategy (Antognazza et al. 2019; Mora et al. 2019; Leempoel et al. 2020) and to detect elusive, rare, and cryptic species effectively even in low density occurrences (Franklin et al. 2019; Shelton et al. 2019; Takahara et al. 2020). Single eDNA sampling could simultaneously monitor biodiversity in the given environment over a broad taxonomic spectrum (Sawaya et al. 2019; Thomsen and Sigsgaard 2019; Zhang et al. 2020). Large DNA barcode reference library that contains barcodes for a wide range of species is essential for successful monitoring of any environments using eDNA. For example; while monitoring marine ecosystems, a previous study could not assign more than 92 percent of the PCR amplified eDNA sequences to any known phyla and the sequences were classified as unassigned phyla (Jeunen et al., 2019; Sawaya et al., 2019). A comprehensive, parameterized reference library with barcodes of most of the locally occurring species and its image data is critical for the reliable application of eDNA technology, and the success of such efforts has been witnessed in probing marine fish diversity (Stoeckle et al., 2020). Barcode of Life Database (BOLD) (www.boldsystems.org) were created with the objective of fulfilling above said requirements (Ratnasingham and Hebert, 2007).

Since only 15 percent of animal species have been barcoded and available in reference libraries (Kvist, 2013), the purpose of this study is to classify amphipods that occur in Vellar estuary sediments using DNA barcoding. We have made considerable efforts in the past decade as part of the Indian Census of Marine Life (ICoML) to recover barcodes in reasonable numbers of marine phyla including fin and shell fishes, invertebrates (Khan et al., 2010, 2011; PrasannaKumar et al., 2012; Thirumaraiselvi et al., 2015; Rajthilak et al., 2015; Rahman et al., 2013, Hemalatha et al., 2016; Palanisamy et al., 2020; PrasannaKumar et al., 2020a, b; Manikantan et al., 2020; Thangaraj et al., 2020) and plants (Sahu et al., 2016; Prasanthi et al., 2020) occurring in and around the Vellar estuary, besides amphipods. This study also aims to test the efficacy of current DNA barcode reference libraries in identifying DNA barcodes, so as to implicitly explain the capacity of the reference library to monitoring environmental quality using eDNA barcodes of amphipods.

## 2. Materials and methods

### 2.1. Sample collection and identification

During February 2012, sediment samples were obtained from the mangroves beds in the Vellar estuary (Latitude: 11° 29’N. Longitude: 79° 46’E.), Southeast coast of India. A total of 6 samples were sampled at multiple sites around the mangrove species; *Rhizopora annamalayana* (Seetharaman and Kandasamy, 2011) using a sterile plastic spatula within the sediment area of 50cm^2^ quadrat. The seaward salinity was 30ppt (measured using hand-held Brix refractometer). The sediment samples were passed with copious ambient seawater through a sieve of 0.5mm pore size and sewn at the site. The amphipods and other fauna were preserved in 95 percent molecular grade ethanol (Merck, India) along with residual sediments and transported to the laboratory. Whenever required, duplicate specimens were preserved for microscope analysis in 5 to 7 percent formaldehyde containing Rose Bengal. Using a Nikon Eclipse E200 compound microscope, the amphipods were sorted and classified to the lowest possible taxonomic ranking based on their morphological characters. For the identification of the specimens, the taxonomic keys of Vinogradov et al. (1996), Martin and Davis, (2001) Bousfield (1978), Balasubrahmaniyan and Srinivasan (1987), Lyla et al. (1999) and Lowry & Myers (2017) which are publicly accessible via the World Amphipoda Database (Horton et al., 2019). Recommended data categories such as geographical co-ordinates, collector, and collection date, unique identifier for voucher specimens, etc., to meet basic data standards for creation of collection database (Evans & Paulay, 2012) was followed. Under the project “DNA barcoding marine amphipods” (tag; DBMA), the barcode data along with image and other meta-data of the present study was released for publically access in BOLD (www.boldsystem.org).

### 2.2. DNA isolation, PCR and sequencing

The DNA was extracted using the DNeasy Blood & Tissue Kits (Qiagen) following the manufacturer’s protocols with 1/10th of actual volume of reagents being adjusted in use. Two or three pereopods were used for DNA extraction, when the individual amphipods were >12 mm in length, and the entire amphipod specimens when <12 mm in length. During the DNA extraction process, elution buffer provided with the kit was used as negative control. Using the primer pair; LCO1490 and HCO2198 (Folmer et al., 1994), the mitochondrial cytochrome c oxidase subunit I (COI) gene was amplified (approximately, 658 base pair). Polymerase chain reaction (PCR) was conducted using a 25μl reaction volume; 12.5μl Taq PCR Master Mix (Invitrogen, India), 11μl distilled water, 0.5μl forward (10 μM) & 0.5μl reverse primer (10 μM), and 0.5μl extracted DNA (50–80 ng/μl). PCR conditions were; initial 2 minute denaturation at 95 °C, followed by 5 cycles at 94 °C at 30 s, 46 °C at 45 s, 72 °C at 45 s and 35 cycles at 94 °C at 30 s, 51°C at 45 s, 72 °C at 45 s, and final elongation at 72 °C at 5 minutes. The negative control processed during DNA extraction used as a negative PCR control. Following PCR, the products were tested on a 1.5 percent agarose gel and commercially Sanger sequenced (bi-directionally) at Macrogen (Seoul, South Korea).

### 2.3. DNA sequence analysis

Sequencing efforts were repeated until at least one specimen in every species captured was sequenced. The sequences were read using ChromasLite ver.2.1 and manually double checked. DNA gaps were tested in BioEdit ver. 7.9 (Hall, 1999) by converting DNA sequences into putative amino acid sequences and aligned in Clustal X ver. 2.0.6 (Thompson, 1997). Properly aligned sequences were submitted to GenBank and made available for public access through accession numbers MT184213-MT184234. Under the project title “DNA Barcoding Marine Amphipods” or by using the tag ‘DBMA’ in BOLD (http://www.boldsystems.org/), meta-data containing voucher information, taxonomy, specimen description and collection data (sample ID, collection date, geographical information, etc.) could be accessed. One could access the dataset provided in the present study at http://dx.doi.org/10.5883/DS-MAOI.

The Barcode of Life Data Systems (BOLD) (Ratnasingham and Hebert, 2007) and GenBank (Benson et al., 2018) were used as a reference library to identify the barcode sequences created in the present study. The comparison of COI sequences in BOLD was facilitated through ‘identification engine’ tool (http://www.boldsystems.org/index.php/IDS_OpenIdEngine) and in GenBank through Basic Local Alignment Searching Tool (BLAST) (Altschul et al., 1990) using a standard similarity search protocol (Hu and Kurgan, 2018). Molecular Evolutionary Genetic analysis (MEGA) version X (Kumar et al., 2018) was used for neighbour-joining (NJ) tree construction using Kimura 2 parameters (K2P). Calculations of the pair-wise distance were rendered using the K2P distance model (Kimura, 1980), nucleotide diversity and Tajima’s statistical test (Tajima, 1989; Nei and Kumar, 2000) were carried out in MEGA X. All 3 codon positions were included in the analysis. For tree based identification, NJ tree was redrawn for better representation in Interactive Tree Of Life (iTOL) (Letunic and Bork, 2019).

## 3. Result and discussion

### 3.1. Species composition and history of its occurrences

We retrieved a total of 2869 amphipod individuals with intact morphometric characteristics. Morphological descriptions assigned the entire collection to 22 species (Fig. 1), 17 genera, and 13 families under the order Amphipoda. List of species described was given in table 1. The Isaeidae family contributed the maximum number of species (n=3), followed by 2 species each in Ampeliscidae, Ampithoidae, Aoridae, Corophiidae, Gammaridae, Maeridae and Talitridae (**Table S1**). The families such as Deaxaminidae, Eriopisidae, Hyalidae, Melitidae and Photidae has contributed single species in the collection.

**Table 1:**
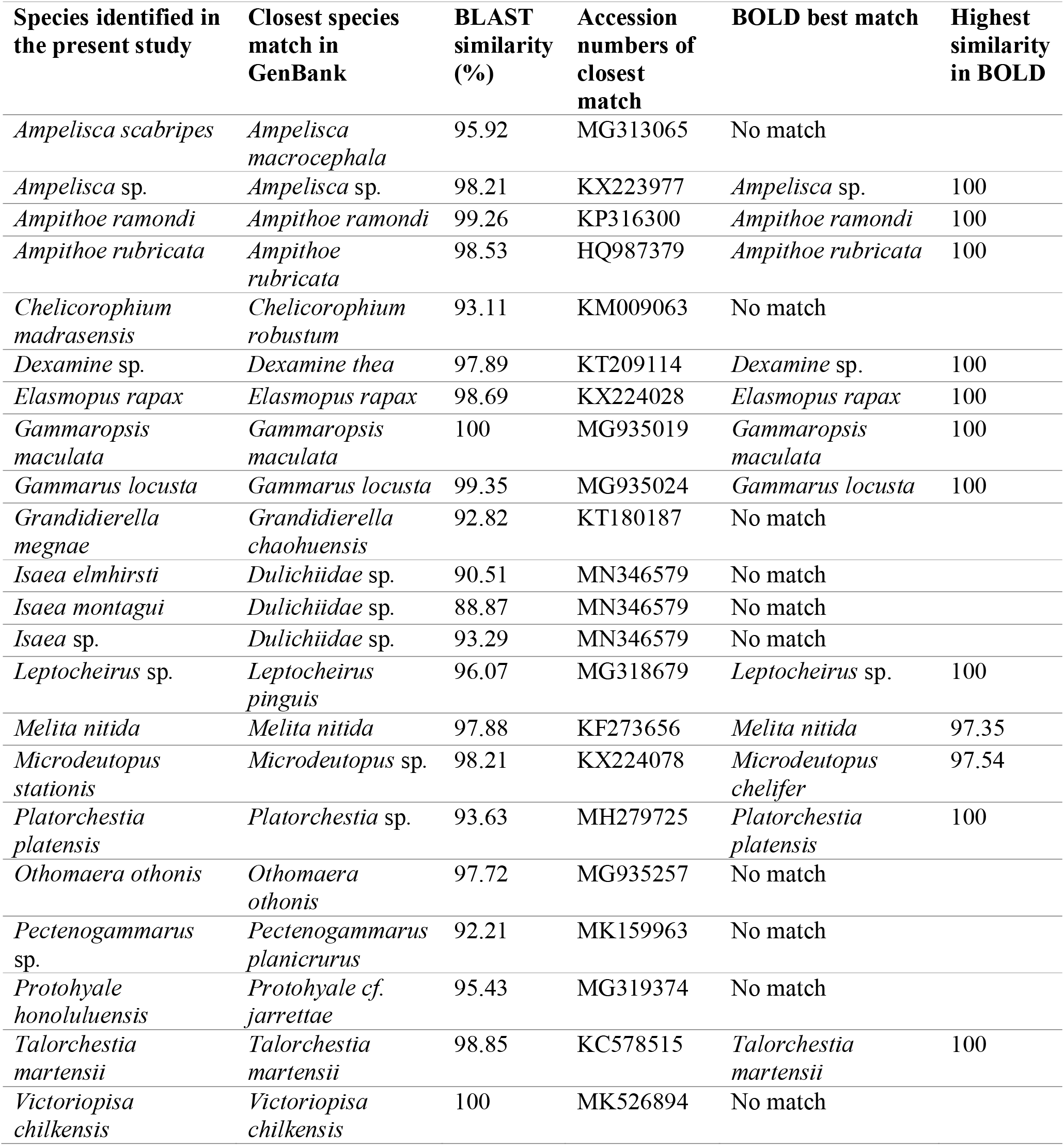
Identification of amphipod COI sequences using GenBank and BOLD databases

| Species identified in the present study | Closest species match in GenBank | BLAST similarity (%) | Accession numbers of closest match | BOLD best match | Highest similarity in BOLD |
| --- | --- | --- | --- | --- | --- |
| *Ampelisca scabripes* | *Ampelisca macrocephala* | 95.92 | MG313065 | No match |  |
| *Ampelisca* sp. | *Ampelisca* sp. | 98.21 | KX223977 | *Ampelisca* sp. | 100 |
| *Ampithoe ramondi* | *Ampithoe ramondi* | 99.26 | KP316300 | *Ampithoe ramondi* | 100 |
| *Ampithoe rubricata* | *Ampithoe rubricata* | 98.53 | HQ987379 | *Ampithoe rubricata* | 100 |
| *Chelicerophium madrasensis* | *Chelicerophium robustum* | 93.11 | KM009063 | No match |  |
| *Dexamine* sp. | *Dexamine thea* | 97.89 | KT209114 | *Dexamine* sp. | 100 |
| *Elasmopus rapax* | *Elasmopus rapax* | 98.69 | KX224028 | *Elasmopus rapax* | 100 |
| *Gammaropsis maculata* | *Gammaropsis maculata* | 100 | MG935019 | *Gammaropsis maculata* | 100 |
| *Gammarus locusta* | *Gammarus locusta* | 99.35 | MG935024 | *Gammarus locusta* | 100 |
| *Grandidierella megnae* | *Grandidierella chaohuensis* | 92.82 | KT180187 | No match |  |
| *Isaea elmhirsti* | *Dulichidae* sp. | 90.51 | MN346579 | No match |  |
| *Isaea montagui* | *Dulichidae* sp. | 88.87 | MN346579 | No match |  |
| *Isaea* sp. | *Dulichidae* sp. | 93.29 | MN346579 | No match |  |
| *Leptocheirus* sp. | *Leptocheirus pinguis* | 96.07 | MG318679 | *Leptocheirus* sp. | 100 |
| *Melita nitida* | *Melita nitida* | 97.88 | KF273656 | *Melita nitida* | 97.35 |
| *Microdeutopus stationis* | *Microdeutopus* sp. | 98.21 | KX224078 | *Microdeutopus chelifer* | 97.54 |
| *Platorchestia platensis* | *Platorchestia* sp. | 93.63 | MH279725 | *Platorchestia platensis* | 100 |
| *Othomaera othonis* | *Othomaera othonis* | 97.72 | MG935257 | No match |  |
| *Pectenogammarus* sp. | *Pectenogammarus planicrurus* | 92.21 | MK159963 | No match |  |
| *Protohyale honoluluensis* | *Protohyale* cf. *jarrettae* | 95.43 | MG319374 | No match |  |
| *Talorchestia martensii* | *Talorchestia martensii* | 98.85 | KC578515 | *Talorchestia martensii* | 100 |
| *Victoriopisa chilensis* | *Victoriopisa chilensis* | 100 | MK526894 | No match |  |

**Fig. 1:**
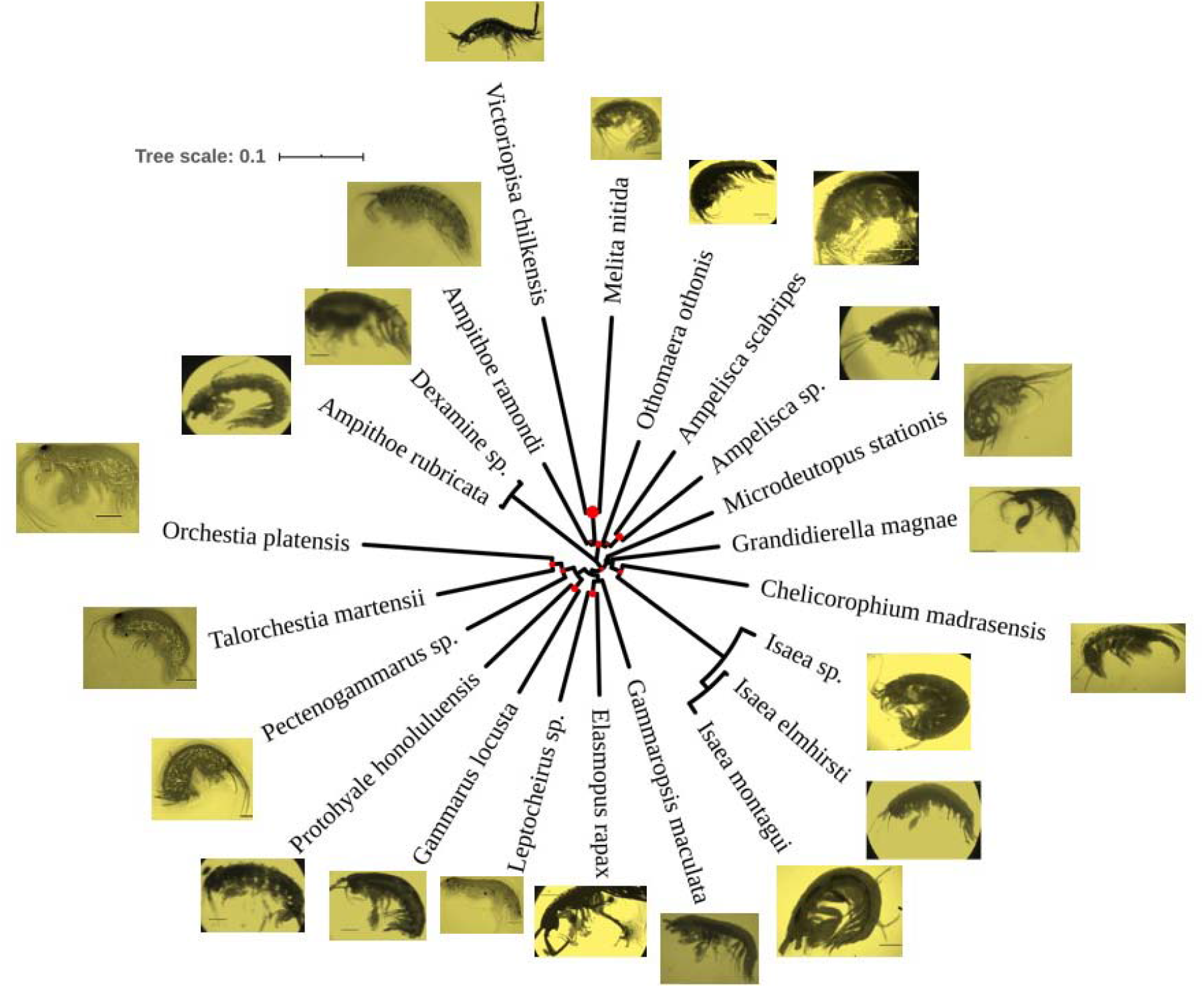
Circular NJ tree drawn using Kimura-2 parametric distance model employing the COI sequences represented by each species of amphipods barcoded. The images represent the duplicate specimens used for image analysis.

Folmer’s primer (Folmer et al., 1994) effectively amplified all 22 species, eliminating the need for additional prime pairs such as those needed for barcoding Atlantic (Costa et al., 2009) and deep sea Pacific amphipods (Jażdżewska and Mamos, 2019). All sequences have been positively confirmed as fragments of the Amphipoda COI gene through BLAST searches. Of the 22 species barcoded in this study, only 4 species *viz., Ampelisca scabripes* (Walker, 1904), *Grandidierella* sp. (Coutière, 1904), *Orchestia* sp. (Leach, 1814) and *Talorchestia* sp. (Dana, 1852) was reported previously in Vellar estuary (Mondal et al., 2010). For the sampled area, more than 81 percent of the species were new records. Further investigation could be directed towards reasoning these new occurrences in Vellar mangrove environment. Hence we investigated previously recorded known habitats of these amphipod species, to perceive any recent notable environmental changes.

Since 1975 (Rabindranath, 1975) till the date (Srinivas, 2019), *Ampelisca scabripes* (Walker, 1904) has been known to occur in Indian estuarine environment. While *Ampithoe ramondi* (Audouin, 1826) occurrences in South Pacific islands were recorded as early as 1986 (Myers, 1986) and their active feeding on leaves and seeds of seagrasses were well known (Castejón-Silvo et al., 2019), their occurrences in the present study are not surprising, given the presence of seagrass patches in Vellar estuary (Ranjitham et al., 2008). *Ampithoe rubricata* (Montagu, 1808) are the most common amphipods previously recorded in kelp forest habitats (Norderhaug et al., 2003), an active red algae feeders (Norderhaug, 2004).

*Chelicorophium madrasensis* (Nayar, 1950) is the continuous feeder recently recorded to dominate amphipod composition of the Cochin estuary sediments along India’s southwestern coast (Rehitha et al., 2019). They have also been recorded elsewhere as a common inhabitant of mangrove forests (Wongkamhaeng et al., 2015). The barcoded *Elasmopus rapax* (Costa, 1853) in this study was previously identf2ied from the coast of Venezuela (Zanders and Rojas, 1992) and its invasiveness is realized in Australian waters (Hughes and Lowry, 2010). Possible invasiveness of this species in mangrove surroundings in Vellar can warrant more investigations. The barcoded *Gammaropsis maculata* (Johnston, 1828) in this study is a good indicator of the complex hydrological forces in an ecosystem 2(Conradi, 2001) and is known to occur in the Tunisian coast of North Africa (Zakhama-Sraieb, 2017). The currently barcoded *Gammarus locusta* (Linnaeus, 1758) was noted for its cosmopolitan estuarine distribution and is expected to be more resistant ocean acidifications (Hauton, 2009). *Grandidierella megnae* (Giles, 1890) barcoded in this study was previously known to occur along Iraq’s Basrah coast (Naser et al., 2010) and on Thailand’s Songkhla Lake (Rattanama et al., 2010). *Isaea montagui* (H. Milne Edwards, 1830) was noted for 2its epibiotic link to crabs (where the amphipod obtains its food from detritus and crab faeces) (Parapar, 1997).

The effectiveness of amphipod DNA barcodes in identifying invasive and delineating cryptic species has been well established (Witt et al., 2006; Bradford et al., 2010; Ros et al., 2015; Lipinskaya et al., 2018). Barcoded in the present study, *Melita nitida* (Smith, 1873) was confirmed to be an invasive species for the Western Scheldt estuary (in Netherlands) that was possibly transported by shipping (Faasse and van Moorsel, 2003). By forming irregular brood plate setae in their bodies (Borowsky et al., 1997), they were known bio-indicators of toxic and petroleum contaminants in sediments. Biomass of molluscan species of Vellar estuary were known to contain significant concentration of petroleum hydrocarbons (Veerasingam et al., 2011). *Microdeutopus stationis* (Della Valle, 1893) barcoded in this study is known to occur in the Tunisian coast of North Africa (Zakhama-Sraieb, 2017) and is abundantly reported in Isles of Scilly seagrass beds (Bowden, 2001).

Herbivorous nature of *M. stationis* may be the explanation for its preference of Vellar mangrove habitat. The barcoded *Platorchestia platensis* (Krøyer, 1845) was previously reported from the Swedish coast and the Baltic Sea (Persson, 2001). *P. platensis*’s ability to reproduce in the warm temperature waters across the South African estuary was argued as one of its characteristics for its invasiveness (Hodgson et al., 2014). *Othomaera othonis* (H. Milne Edwards, 1830) barcoded in this study is known to occur in northern Atlantic Ocean in the Portuguese continental shelf (Sampaio et al., 2016) and in the Algerian, Mediterranean Sea continental shelf (Bakalem et al., 2020). However its occurrences in shallow mangrove sediments were unknown until now and requires further investigation. The barcoded *Protohyale honoluluensis* (Schellenberg, 1938) in this study was known to occur in the Hong Kong Island marine caves of (Horton, 2008).

The barcoded *Talorchestia martensii* (Weber, 1892) in this study is generally referred to as equatorial sandhoppers occurring in African beach sand (Ugolini, 2016) and Kenyan coast (Ugolini and Ciofini, 2015). *T. martensii* was widely used astronomical orientations researches (Ugolini and Ciofini, 2015; Ugolini, 2016). *Victoriopisa chilkensis* (Chilton, 1921) barcoded in the present study was known to occur on Malaysian coast (South China Sea) and was used as feed in Thailand’s shrimp cultures (Yokoyma et al., 2002). Neither traditional nor molecular approaches could resolve the four taxa viz., *Ampelisca* sp., *Dexamine* sp., *Isaea* sp., *Leptocheirus* sp., and *Pectenogammarus* sp. recorded in this study to species level identification. In near future, these barcodes in the public databases may be resolved to species level when the respective barcodes of the species were obtained elsewhere.

### 3.2. Sequence analysis and species identification

The COI sequences represented by the families viz., Ampeliscidae [*Ampelisca* sp. (Krøyer, 1842), *A. scabripes* (Walker, 1904)], Corophiidae [*Chelicorophium madrasensis* (Nayar, 1950), *Leptocheirus* sp. (Zaddach, 1844)], Aoridae [*Grandidierella megnae* (Coutière, 1904)], Talitridae [*Platorchestia platensis* (Krøyer, 1845), *Talorchestia martensii* (Weber, 1892)), Hyalidae (*Protohyale honoluluensis* (Schellenberg, 1938)] and Isaeidae [*Isaea* sp. (H. Milne Edwards, 1830), *I. elmhirsti* (Patience, 1909) and *I. montagui* (H. Milne Edwards, 1830)] were barcoded for the first time (59% of the sequences). Over 59% of the barcoded species in this study was missing in the reference libraries. The COI sequences of the Isaeidae family produced in the present study were found to be sequenced for the first time, because previously no COI barcodes of this family were found in the GenBank library. Even when the COI barcodes where referred across in the BOLD database, BOLD declared the first time barcoded species as “no match” in their database (Table 1). In general, significant amounts of barcode records were resolved up to family level in the reference database, partly because 90% of multicellular organisms remain undescribed (Kwong et al., 2012; Curry et al., 2018) and the Vellar estuary could be hot spot for marine fauna and flora yet to be barcoded (Khan et al., 2010; PrasannaKumar et al., 2020a, c; Manikantan et al., 2020; Prasanthi et al., 2020).

Since 18 species among the 22 barcoded species were not recognized to occur during previous study (Mondal et al., 2010), the development of this local barcode library would help in the near future to monitor amphipod species diversity and composition through eDNA barcoding surveys. As seen in few marine amphipod species of Canadian waters (Tempestini et al., 2018), we detected no stop codons in the COI sequences. Mean GC content was 39.25±0.93% which is slightly higher than the Arctic amphipod’s GC content of (32.9%) (Tempestini et al., 2018). Table 2 provides descriptive statistics for the distribution of nucleotide frequencies.

**Table 2:** Summary statistics for nucleotide frequency distribution.

| Nucleotide composition (%) | Minimum | Mean | Maximum | Standard error |
| --- | --- | --- | --- | --- |
| Guanine (G) | 12.89 | 19.83 | 26.92 | 0.7789 |
| Cytosine (C) | 14.19 | 19.69 | 26.1 | 0.5526 |
| Adenine (A) | 21.21 | 25.46 | 31 | 0.5843 |
| Thymidine (T) | 29.85 | 35.03 | 39.8 | 0.5434 |
| GC (all codon positions) | 31.65 | 39.52 | 46.82 | 0.9315 |
| GC-Codon Position 1 | 37.25 | 45.83 | 51.96 | 0.8605 |
| GC-Codon Position 2 | 39.02 | 42.22 | 43.9 | 0.302 |
| GC-Codon Position 3 | 13.24 | 30.48 | 45.59 | 2.0274 |

Of the 22 species barcoded, least inter-specific pair-wise distance of 0.16% was observed between *Isaea elmhirsti* and *I. montagui* (**Table S2**). The maximum inter-specific distance of 5.51% was observed between *Victoriopisa chilkensis* and *Orchestia platensis* (**Table S2**). The inter-specific maximum (0.16%) and minimum (5.51%) values observed were lower than what those reported in Canadian amphipods (0.6% and 18.07%, respectively) (Tempestini et al., 2018). The interspecific values observed for deep-sea amphipod species (13%) (Knox et al., 2012) was also higher than the maximum interspecific values observed in this study. Although a 16% threshold has been suggested for the delineation of amphipod species (in Gammaridae or Niphargidae) (Lefébure et al., 2006; Flot et al., 2010; King et al., 2011; Fiser et al., 2015), lower values have also been recorded for Talitridae (8 - 17%) (King et al., 2011) or for Hyalleidae (4%) (Witt et al., 2008).

When we averaged the pairwise distance of each species to the rest of the species barcoded in this study, we found *Melita nitida* and *Victoriopisa chilkensis* had the maximum pairwise average distance (0.45) (Fig. 2). By including COI sequences from 13 families, we show that the threshold of 16% was too high to delineating most of the species barcoded in this study and proposed inter-specific values of 26.8% for Gammaridae species (Costa et al., 2009) was more than 4 times the maximum inter-specific values observed among the species in this study. Similar observations were also reported in Canadian amphipods (Tempestini et al., 2018). The interspecific divergence among the species of *Gammarus* genera was found to be more than 20% (Hou et al., 2009).

**Fig. 2:**
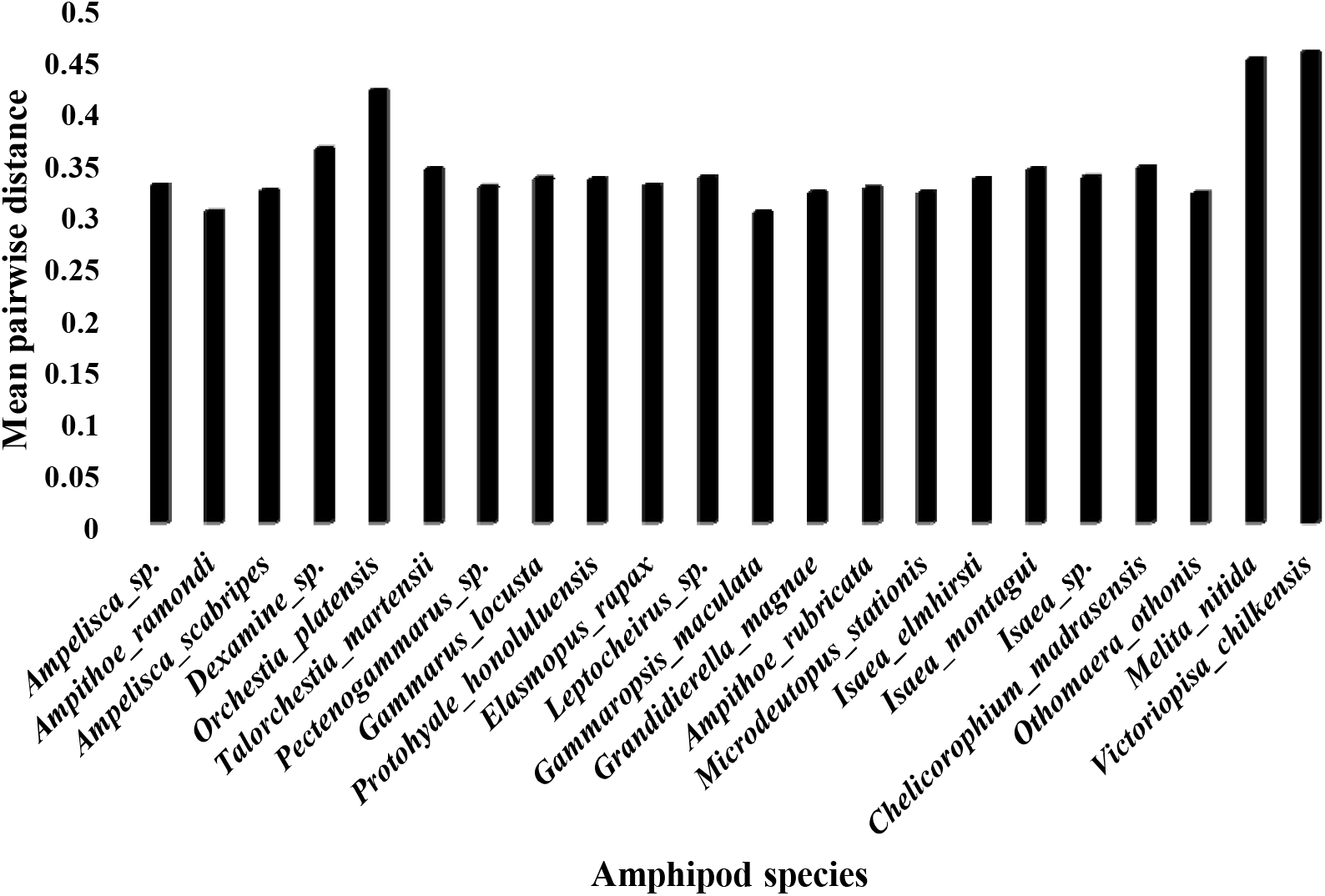
Average pair-wise distance between each species versus rest of the amphipod species barcoded.

Hence, it is becoming clearer that each amphipod family have differential species threshold as genetic variability among amohipods depends on family traits such as habitat preferences and geological history (Knox et al., 2020). At the same time family traits alone cannot be reasoned, as there exists a contrasting genetic structure even among closely related amphipod species (Baird et al., 2011). Hence defining family specific species threshold for delineating amphipod species is necessary.

The overall average pair-wise distance was 3.59%. Nucleotide diversity and Tajima’s statistics were 0.27 and 2.62, respectively. We found that 34.86% were conservative sites and 58.2% was parsimonious from the final alignment.

Though the percentage of conservative sites were slightly higher than Canadian amphipods (23.5%), percentage of parsimonious sites were slightly lower when compared to Canadian amphipods (69.25%) (Tempestini et al., 2018). Using a single gene for species identification may introduce bias in different steps of sample processing (Radulovici et al., 2010), lack of established barcode gap (Meier et al., 2008), and superficial increase in species threshold estimates by ‘COI-like’ sequences (Buhay, 2009). Hence we also conducted tree-based species identification using clustering analysis.

### 3.3. Tree based identification

Based on the statistical significance (percentage of identity, query coverage, e-value), reference sequences for tree based identifications were retrieved from GenBank. Details of all sequences obtained during BLAST analysis was listed in **Table S3**. The first time sequenced species which did not had significant match in the database were used as such in the construction of the NJ tree without any reference sequence (Fig. 2). All COI sequences generated in this study (n=22), clustered with the reference sequences (n=24) of respective species in the same branch (Fig. 3). Most of the branches (>90%) in the NJ tree were backed by maximum (>75) bootstrap values. The references sequences precisely clustered with its corresponding species barcoded in this study, suggesting the success of tree based identification. The NJ tree constructed using COI sequences generated 3 major clades (Fig. 3). A minor clade consisting of two genera (*M. nitida* and *V. chilkensis*) was the first and top most clade in the tree. There are 3 genera (*Ampithoe, Ampelisca* and *Othomaera*) in the second and middle clade.

**Fig. 3:**
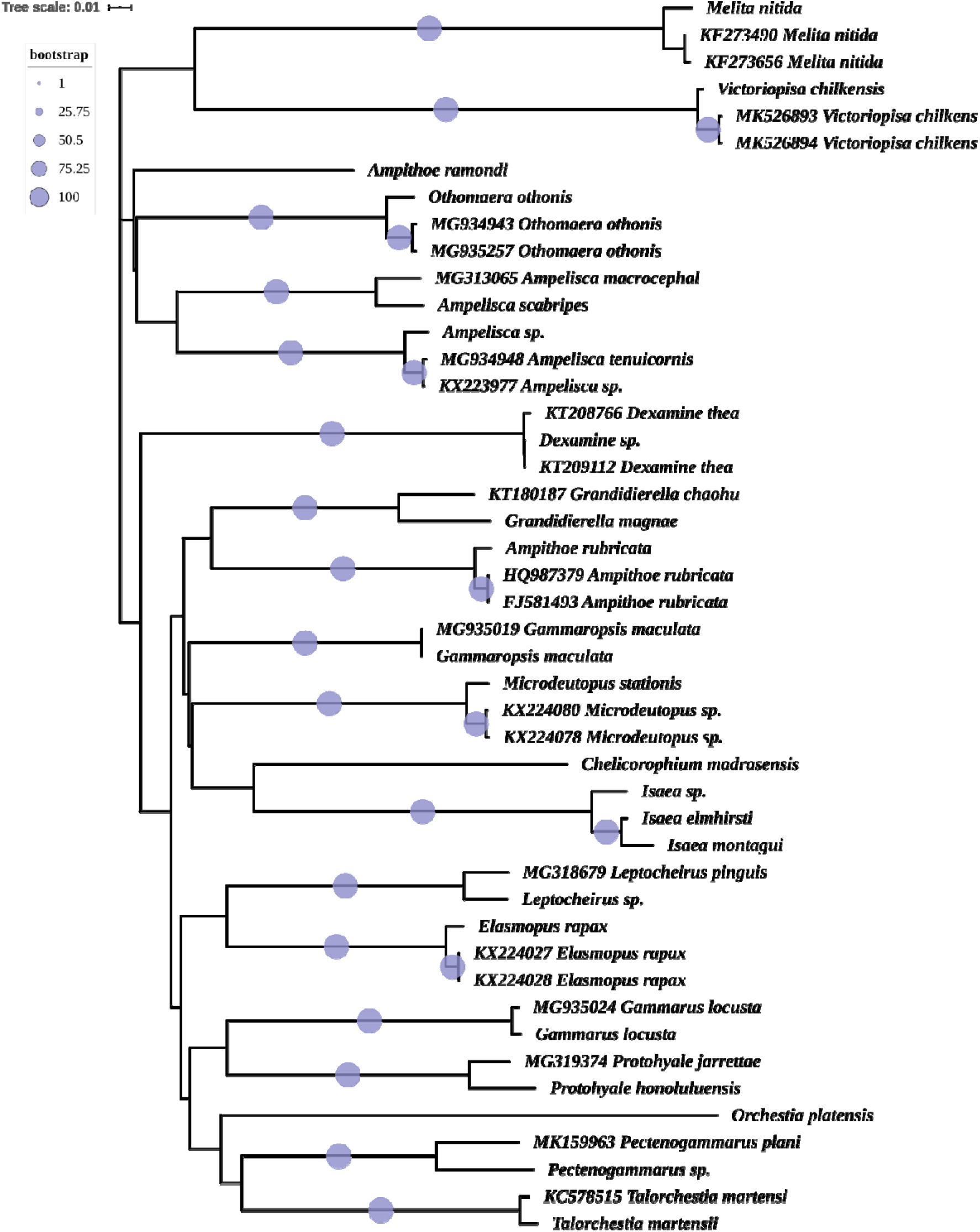
NJ tree was drawn using COI sequences with Kimura-2 parametric distance model. The sequences retrieved from GenBank were represented by “accession-number_species-name”. Example; “MG935024_*Gammarus locusta*”. The sequences of the present study were represented with species name only. Time scale and bootstrap legends were given at the top left corner of the tree. More than 90% of the branches were supported by >75% of bootstraps values.

The largest clade consisting of 14 genera was the bottom-most and the third clade. 3 subclades were evident within the third clade, where the genera *Dexamine* constituted the smallest sub-clade. The second sub-clade contained the members of Ampithoidae, Aoridae, Corophiidae, Isaeidae and Photidae. The bottom-most and third sub-clade contained the members of Corophiidae, Gammaridae, Hyalidae, Maeridae and Talitridae. While certain members of the same family (example; Gammaridae, Corophiidae) or the same genera (example; *Ampithoe*) did not cluster together, sequences of same taxa cluster together suggest the efficacy of COI sequences in species delineation. There was also evidence that the members of the same family (example, Isaeidae) forming single cluster.

### 3.4. Environmental monitoring

Amphipods are commonly used to assess sediment quality in marine and estuarine environments (Chapman 1992, 2013; Postma et al., 2002). High throughput sequencing has enabled the identification of array of species present in the environment by means of its eDNA presence in the environmental substances (Cristescu, 2014; Deiner et al., 2014; Braukmann et al., 2019). This eDNA barcoding approach reveals species composition of entire trophic levels, from complex bulk environmental samples, gut or faeces content in soil and in aquatic environments or from ancient samples collected within permafrost or in sediments (Cristescu, 2014; Deiner et al., 2017). Thus the composition of amphipod species revealed by such approach will in turn disclose the environmental quality (polluted, nonpolluted, toxic compounds, etc.). The eDNA barcoding begins by collecting a large number of sequences from one or more standard barcode region targeting a specific groups or taxa (Hebert et al., 2018), followed by comparing these sequences with the reference sequences in the databases like GenBank and BOLD.

More species coverage in the reference database is strongly required, as the sequencing costs of next generation sequencing are reducing (Hebert et al., 2003; Shokralla et al., 2014; Cruaud et al., 2017) and sequences from museum and type specimens could be obtained (Prosser et al., 2016). Since 59% of the species barcoded in this study were first time sequenced and released for public access, the present data set will aid to improve species coverage in the databases and encourage better monitoring of the environment. The reliability of environmental and meta-DNA barcoding studies depends on validated reference library generated through taxonomic expertise, accessible image data, voucher specimens and their genetic materials. The dataset developed through this study has been made available in both GanBank and BOLD. For the present study dataset, a unique digital object identifier has been created in BOLD, which can be accessed at http://dx.doi.org/10.5883/DS-MAOI. The voucher specimens were accessible through the Centre Advanced Study in Marine Biology, Annamalai University’s Marine Museum, which will allow future referencing analysis or avoid identification error cascades (Bortolus, 2008). Cryopreserved genetic materials were also accessible upon request, as we realised the values of such materials in construction of reference library (Hanner and Gregory, 2007; Gonzalez et al., 2018).

## 4. Conclusion

We show that, reference libraries (GenBank and BOLD) have to be strengthened for improved barcode based amphipod species identification, as small sequencing effort such as ours discovered 13 species as first time barcoded. While our data set represents local custom library for the identification of amphipod species with limited taxonomic reach (n=22 species), with meta-data ready for accession, they have become less prone to errors (Machida et al., 2017; Heller et al., 2018). The current study provides a valuable reference library particularly for those species that were first time barcoded, against which barcodes of marine amphipods from different regions can be referred in the near future, as these parameterized reference libraries will be crucial to species identification (Ekram et al., 2007; Wilson et al., 2011). Since amphipods are widely used as a universal taxonomic screening tool in environmental monitoring, amphipod barcodes along geographic and ecological data, may not only promote our knowledge on taxonomy, phylogeography, and species crypticism, but also serve as a powerful tool for environmental monitoring and health assessment. It should be noted that the DNA barcoding technology is growing beyond systematic or taxonomic study. At the same time, the developing high-throughput sequencing technologies massively alters environmental surveys and bio-monitoring applications (Fonseca et al., 2010; Hajibabaei et al., 2011; Leray et al., 2015). As a result, reference datasets such as ours will become important for health assessment and environmental monitoring using amphipod barcodes. Although the creation of a comprehensive amphipod barcode library with extensive species coverage was an ultimate goal (example; Zahiri et al., 2017) this study represents a small step towards it.

## Acknowledgement

First author thanks the INSPIRE Fellowship of the Department of Science and Technology (IF10431), India and China Postdoctoral Research Foundation (0050-K83008), China for their financial assistance. We thank Raja S for his assistance with the taxonomic knowledge and the verification of species. We acknowledge Ms. Clare Adams, Department of Anatomy, Dunedin, New Zealand’s efforts for critically reviewing the manuscript and improving its content. We thank the anonymous reviewers for improving the quality of the manuscript.

